# DeltaAI: Semi-Autonomous Tissue Grossing Measurements and Recommendations using Neural Radiance Fields for Rapid, Complete Intraoperative Histological Assessment of Tumor Margins

**DOI:** 10.1101/2023.08.07.552349

**Authors:** Anish Suvarna, Ram Vempati, Rachael Chacko, Gokul Srinivasan, Yunrui Lu, Brady Hunt, Veronica Torres, Kimberly Samkoe, Matthew Davis, Lucy Fu, Brock Christensen, Louis Vaickus, Matthew LeBoeuf, Joshua Levy

**Affiliations:** Thomas Jefferson School for Science and Technology, Alexandria, VA; Geisel School of Medicine, Hanover, NH; Department of Pathology and Laboratory Medicine, Dartmouth-Hitchcock Medical Center, Lebanon, NH; Department of Medicine, Section of Radiation Oncology, Dartmouth-Hitchcock Medical Center, Lebanon, NH; Thayer School of Engineering, Dartmouth College, Hanover, NH; Department of Dermatology, Dartmouth-Hitchcock Medical Center, Lebanon, NH; Dartmouth College, Hanover, NH; Department of Epidemiology, Dartmouth College Geisel School of Medicine, Hanover, NH; Department of Molecular and Systems Biology, Geisel School of Medicine at Dartmouth, Lebanon, NH 03756; Department of Community and Family Medicine, Geisel School of Medicine at Dartmouth, Lebanon, NH 03756; Program in Quantitative Biomedical Science, Dartmouth College Geisel School of Medicine, Hanover, NH

## Abstract

Mohs Micrographic Surgery (MMS) aims to excise cutaneous cancer with real-time margin analysis. However, manual tissue grossing and analysis can be inefficient, so we propose DeltaAI, a novel workflow that utilizes Neural Radiance Fields (NeRF) to enable rapid tissue grossing and generate a 3D model in an augmented reality (AR) environment. In our study, we captured 30-second videos of 17 MMS specimens using a photogrammetry turntable and cellphone camera. Preprocessing the tissues with segmentation models, we created a dataset of 923, 360-degree-view, images per video (17 videos). Using COLMAP, we estimated poses for sparse tissue reconstructions and trained the NeRF model for 3D volumetric tissue renderings. The results demonstrated that DeltaAI generated more accurate and complete 360-degree, 3D tissue renderings compared to previous models, while also achieving significantly faster runtimes. Our proposed semi-autonomous NeRF-based workflow has the potential to enhance the speed of MMS specimen processing, measurement, report generation, and margin assessment. It can inform real-time grossing decisions, automate the export of electronic health record data, and facilitate time-efficient and complete cancer excisions. Moreover, DeltaAI can contribute to the wider adoption of AI technology in clinical settings by improving tissue modeling for manual grossing.

## Introduction

### Background

Surgical resection is the most common method for removing solid cancer tumors. It involves removing the affected tissue along with the surrounding tissue, which is then examined in a laboratory (Chandrasoma, 2018). Intraoperative real-time assessment is often limited due to time and labor constraints. However, wide surgical margins can cause aesthetic and functional compromises, while undersampling of tissue margins can result in incomplete excision and tumor recurrence (Ecclestone et al., 2020). To avoid complications and additional costs, a more thorough study of tissue margins is needed as tumors can extend beyond visible areas (Levy et al., 2022).

### Mohs Micrographic Surgery (MMS)

Mohs Micrographic Surgery (MMS) offers rapid and thorough margin assessment (Mastacouris & Mafee, 2021) during surgery. This technique involves layer-by-layer tissue removal, minimizing tissue loss and enabling comprehensive margin study (Finley, 2003). A specialized surgical team, including a surgeon, histo-technician, and pathologist, performs real-time examinations and communicates findings. A surgical tumor map is created to identify positive margins and guide further removal if needed (Levy et al., 2022). Compared to standard techniques, MMS has significantly improved patient outcomes, with a cancer recurrence rate of less than 2% versus over 20% for standard excision (Bialy et al., 2004).

### Need for Improved Tissue Grossing and Limitations of Prior Art

Tissue grossing is a critical component of surgical pathology, providing vital information for accurate diagnosis and treatment planning. Gross examination of tissue involves macroscopic assessment of size, color, texture, and consistency, as well as the identification of gross abnormalities and suspicious lesions, playing a factor in disease staging. Tissue grossing is also a crucial component of specimen preparation for histological assessment, forming an initial impression of the tissue properties prior to optimal specimen sectioning. In MMS, specimen sectioning is highly specialized, ink is applied to the tissue to define a coordinate system that is correspondent to similar markings on the original surgical site and surgical tumor map. Despite its significance, manually grossing tissues is a time-consuming process with considerable inter- and intra-observer variability [REF]. Furthermore, manual grossing can be prone to errors and can result in incomplete sampling/sectioning or inadequate representation of the tissue specimen, leading to misdiagnosis or suboptimal treatment outcomes.

To address the limitations of manual grossing, there has been growing interest in the development of automated tissue grossing techniques. Automating the reporting of the gross tissue dimensions and making inking and sectioning recommendations based on the tissue characteristics has the potential to boost productivity and efficiency, minimize inter-observer variability, and improve tissue evaluation accuracy and repeatability. Several semi-automated grossing systems which leverage robotics, computer vision, and machine learning have been designed though have not yet been proven to be tractable for incorporation in the intraoperative setting due to insufficiencies in tissue modeling with significant time constraints (Cheng et al., 2021). Semi-autonomous tissue grossing can be enhanced through the generation of 3D models of tissue/tumor, which can provide more comprehensive information about the gross tissue/tumor morphology. This can facilitate preoperative planning, intraoperative grossing, and postoperative evaluation. Incorporation of 3D models into augmented reality and virtual reality computer applications can provide technicians and surgeons with a more natural and immersive approach to analyzing tumors.

### Cell Phone App

The proposed model, DeltaAI, combines Neural Radiance Fields (NeRF) with the ArcticAI workflow to enhance the accuracy and efficiency of Mohs Micrographic Surgery (MMS). DeltaAI generates a 360-degree, 3D tissue model in under 30 seconds, providing precise measurements and orientation indicators for improved gross morphology reporting and tissue sectioning. NeRF fills in gaps in the tissue sample and enhances image quality. With real-time 3D modeling and advanced analytics, DeltaAI offers automated and accurate tissue assessment, reducing the risk of tumor recurrence. Its speed enables faster tissue processing, making it suitable for complex cases and improving patient outcomes. DeltaAI is a key component for future automation in MMS, facilitating real-time margin assessment and streamlining surgical workflows.

Stereology, laser scanning microscopy, and confocal microscopy have previously been used to generate 3D models of solid tumors for assessment of gross morphology (Bisson-Larrivée & LeMoine, 2022). These methods can be time-consuming, technically demanding, and possibly call for expensive equipment or expertise. Other popular techniques include LiDAR scanning (i.e., pulsed lasers to measure ranges) or 3D cameras (i.e., depth sensing through multiple cameras), or in the absence of such technologies, triangulation of similar imaging features across images from multiple viewpoints (the cornerstone of photogrammetry, structure from motion and multi-view stereo techniques).

Deep learning-based segmentation allows for the identification of features in images, such as tissue, while NeRF can generate segmented images from various perspectives quickly. The combined use of deep learning and NeRF is ideal for creating superior 3D tissue models. Challenges remain for their application in 3D modeling of gross tissue specimens for MMS, including: 1) the development of robust and accurate algorithms for tissue segmentation, feature extraction, and model generation, and 2) the integration of automated techniques into the pathology workflow, including sample preparation, imaging, analysis, and reporting. These algorithms may be confounded by variations in tissue type, size, and quality, as well as account for artifacts and distortions introduced during tissue processing and staining.

Current 3D modeling of the gross specimen leverages triangulation techniques to create 3D point clouds, which can introduce gaps and holes for points that are unable to be fully resolved (LeBoeuf et al., 2020) (Levy et al., 2022). The use of triangulation-based techniques to reconstruct point clouds is also likely to be impractical for real-time analysis during surgery due to time inefficiencies introduced through the pairwise comparison of all images extracted during the examination of tissue (Levy et al., 2022). The primary objective of this study was to improve the efficiency of surgical workflows in MMS by automating and standardizing tissue grossing practices and to provide an extended discussion of previously referenced neural network workflows for 3D gross specimen modeling. Optimizing the sectioning and inking of tissue and generating smart grossing recommendations can significantly reduce the time needed to determine if further excision is required. To address the limitations of current 3D specimen grossing we developed a new 3D modeling algorithm called DeltaAI based on NeRF. This algorithm allows for rapid 3D model generation and accurate tissue grossing recommendations and reports. Our system calculates and creates precise measurements and orientation indicators within seconds using NeRF and other Artificial Intelligence algorithms. As a result, our platform eliminates the need for manual measurements and significantly reduces examination time during MMS, while also enabling the exporting of relevant information to EHR.

## Methods

### Data Collection

Photogrammetry techniques were used to create 3D models of tissue samples prior to histological assessment. A cost-effective photogrammetry studio was developed to facilitate the imaging of gross Mohs excisions (all from Basal Cell Carcinoma excisions) at multiple viewing angles. During Mohs surgery, prior to specimen grossing, seventeen large tissue samples were positioned on circular turntables, and video was recorded using a phone camera (iPhone 12; Apple, Cupertino, CA) placed at a fixed distance from a turntable, acquiring nearly one thousand imaging frames of the turntable setup for further analysis. Approximately twenty-four-second videos were recorded for each piece of tissue. The length, width, and height of the tissue dimensions were recorded with the help of surgical pathology staff.

### Tissue Preprocessing

A segmentation model was developed to isolate the tissue from the surrounding background (e.g., turntable) within each imaging frame. We preprocessed the tissue videos in this manner to generate a dataset of 923 images per video (17 total videos), including images sampled from 360-degree viewpoints. Videos were subsampled down to 20-30 frames to simulate data collection using a faster turntable.

### 3D Tissue Modeling

Neural Radiance Fields (NeRF) is a machine learning architecture that adopts a generative modeling approach to gain a 3D comprehension of the gross morphology of the tissue (Müller et al., 2022). It achieves this by training to interpolate images of the tissue (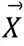) from various camera positions/directions (*x*, *y*, *z*, θ, ϕ; akin to the spatial coordinates, yaw, and pitch– roll does not require additional image manipulation), which allows the synthesis of new viewpoints of the tissue. The primary novelty of the model is the adoption of neural graphics primitives, a sampling technique to drastically lower NeRF’s computational complexity, enabling real-time rendering of 3D scenes on low-cost hardware (e.g., cell phones). The DeltaAI workflow utilizes a NeRF model trained with hash encodings that enable rapid convergence. Its capacity to create intricate 3D models from a small collection of 2D photos motivated the selection of this approach, as input to the model camera intrinsic orthogonal, parallel to, and top-down with respect to the tissue allow for rapid length, width, and height measurements common in the Mohs surgical context. In addition, NeRF models can automatically render high-quality tissue images devoid of gaps/holes in the tissue render, something other techniques are unable to accomplish (Müller et al., 2021). To rapidly estimate the camera positions/directions, we employed COLMAP, a library, and platform created by Johannes L. Schönberger and Jan-Michael Frahm, to estimate the poses (e.g., camera position/orientation– *x*, *y*, *z*, θ, ϕ), serving as the input to NeRF along with each corresponding image (denoting as *DeltaAI*). As a comparison, 3D point clouds were generated from this data using a sparse reconstruction approach (denoting as *SparseCOLMAP*). We further combined multiple sparse 3D point clouds into a dense reconstruction of the tissue (denoting as *DenseCOLMAP*). In this manuscript, the implementation of COLMAP differs from the previously reported workflow by excluding certain pre-/post-processing steps, allowing for a fair and direct comparison between the two workflows with similar pre-processing steps. The performance of this workflow was then compared to MeshLab, an unrestricted and fully-accessible program capable of performing similar photogrammetry techniques, generating a 3D point cloud visualization from a set of 2D images (method denoted as *MeshLab*) (Cignoni et al., 2008). All models were optimized using graphics processing units (GPUs). A calibration technique was developed which used the dimensions of segmented tissue in relation to the turntable to report tissue dimensions (length, width, height) in centimeters.

### Model Comparison

Four approaches (*DeltaAI, COLMAP-Sparse, COLMAP-Dense, MeshLab*) were assessed and compared based on two criteria: 1) computational time to produce 3D tissue models, and 2) agreement of their measures for length, width, and height with ground truth measures.

The mean computational time was estimated and compared between approaches using Bayesian hierarchical log-normal regression modeling. A Bayesian approach to compare times was used due to low sample size– Bayesian methods help avoid bias through the specification of noninformative priors. Using this regression framework, the approach (*DeltaAI, COLMAP-Sparse, COLMAP-Dense, MeshLab*) was modeled as a categorical fixed effect, with time as the dependent variable and specimen as a random effect. This reported mean convergence time along with 95% credible intervals. Statistical significance was assessed using estimated marginal means, which performed pairwise comparisons between algorithmic fit times, reporting 95% credible intervals and p-values (derived from the probability of direction, *pd; p*≈2 * (1 − *pd*)).

Mean absolute measurement error *d* for each of the 3D tissue dimensions (e.g., *d* = |*L_actual_* − *L_predicted_*|) was calculated using log-linked Gamma Bayesian hierarchical regression models, which function similar to the lognormal approach. The absolute difference (i.e., estimating mean absolute error) between the actual and algorithmically derived tissue dimensions served as the dependent variable, while an interaction term between the approach (*DeltaAI, COLMAP-Sparse, COLMAP-Dense, MeshLab*) and each image dimension (length–L, width–W and height–H) allowed for comparisons between the methods by each image dimension:

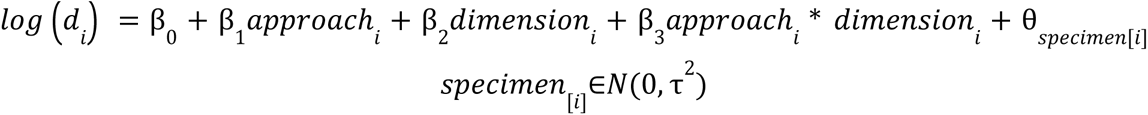

Mean absolute measurement error and statistical significance were reported and compared across methods for each tissue dimension, similar to the previous statistical model used for time. The proportional difference (estimating mean proportional error– e.g., 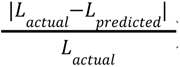) was calculated and compared using a similar approach. The overall performance of *DeltaAI* and *Colmap* approaches were compared to each other, independent of the tissue dimensions using leave-one-out performance metrics, which was used to formulate a statistical test akin to a likelihood ratio test, returning a z-score and p-value indicating the preference of one method over another.

## Results

Using the DeltaAI workflow, only 20-30 images for each sample were needed for accurate estimation of the camera parameters and 3-dimensional reconstruction (**Supplementary Table 1**). The NeRF model was trained on the camera position/orientation along with corresponding images, and taking into account this convergence along with pose estimation– the *DeltaAI* workflow was able to run in under 30 seconds on average. This platform resulted in a thorough reconstruction of the tissue specimen, allowing for the placement of necessary grossing and orientation indicators, with no holes, gaps, or other inconsistencies (**Figure 3**). Importantly the model can: 1) estimate the tissue by directly generating images from the top, side, and front views, 2) place orientation indicators (where to place red/blue inks; used for determining position/orientation of tissue at the surgical site) and 3) assess grossing indicators (e.g. demonstrate where to bisect tissue prior to serial sectioning as shown in **Figure 3**), all of which are critical both for documentation and deciding if/where additional tissue needs to be excised. Videos demonstrating interaction with the 3D tissue specimen and placement of the specimen sectioning and orientation indicators can be found at the following URL: https://deltaai.netlify.app/.

### Runtime Results

DeltaAI was able to 3D reconstruct tissue in just under 30 seconds for both radial and peripheral margin assessment, significantly faster than all other approaches (**Figure 2A**). The second fastest approach (SparseCOLMAP), took approximately three times as long to converge and was significantly slower than DeltaAI (B=-1.19, 95%CI [-1.29– -1.09], p<0.001) (**Table 1**; **Figure 2A**). Important for setting reliable expectations during tissue grossing, the runtime for DeltaAI was also far more consistent than the other approaches (95%CI [27.2-31.8s]).

**Figure 1:**
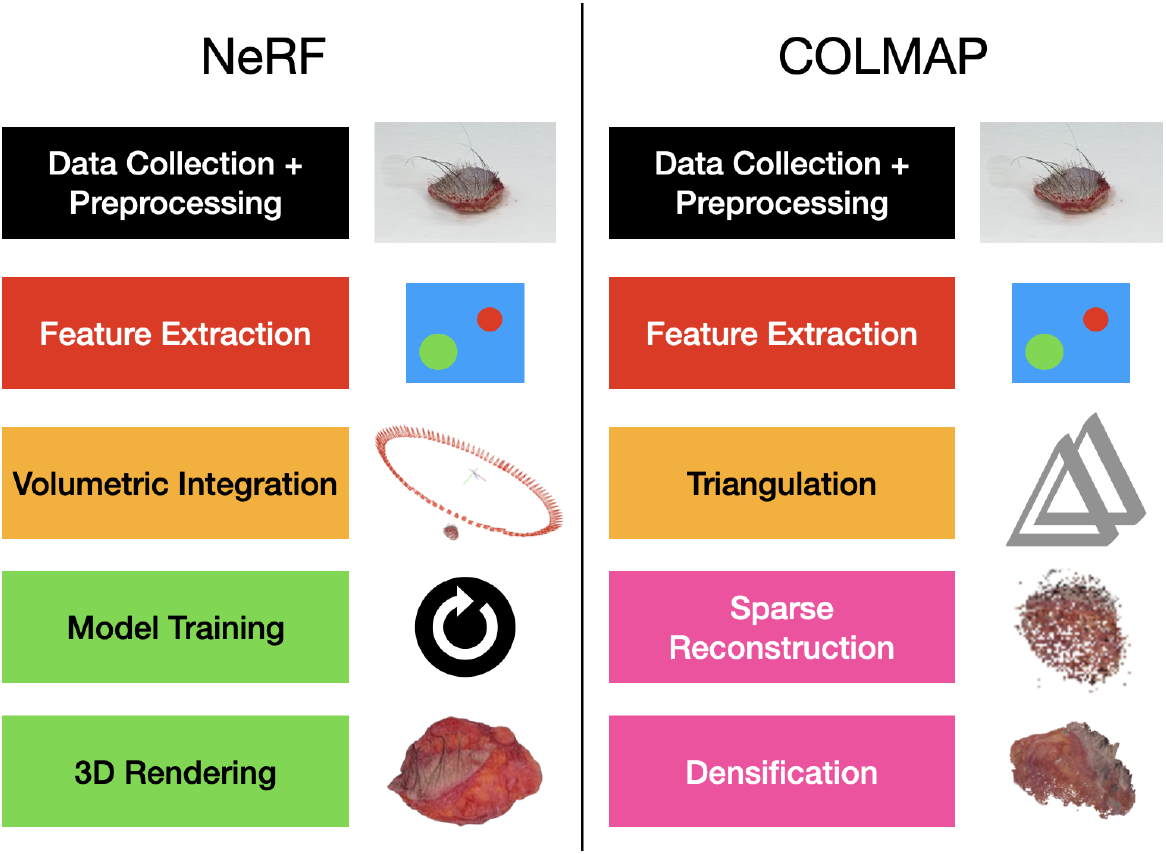
3D Modeling Methods Overview. The left side contains the overview for the Neural Radiance Fields workflow, which is the proposed method of this paper. The right side contains an overview of COLMAP, an existing photogrammetry technique. The data collection, preprocessing, and triangulation/volumetric-integration steps are similar, but the later steps vary between the methods, allowing for NeRF to excel in eliminating inconsistencies and gaps in the 3D model convergence.

**Figure 2:**
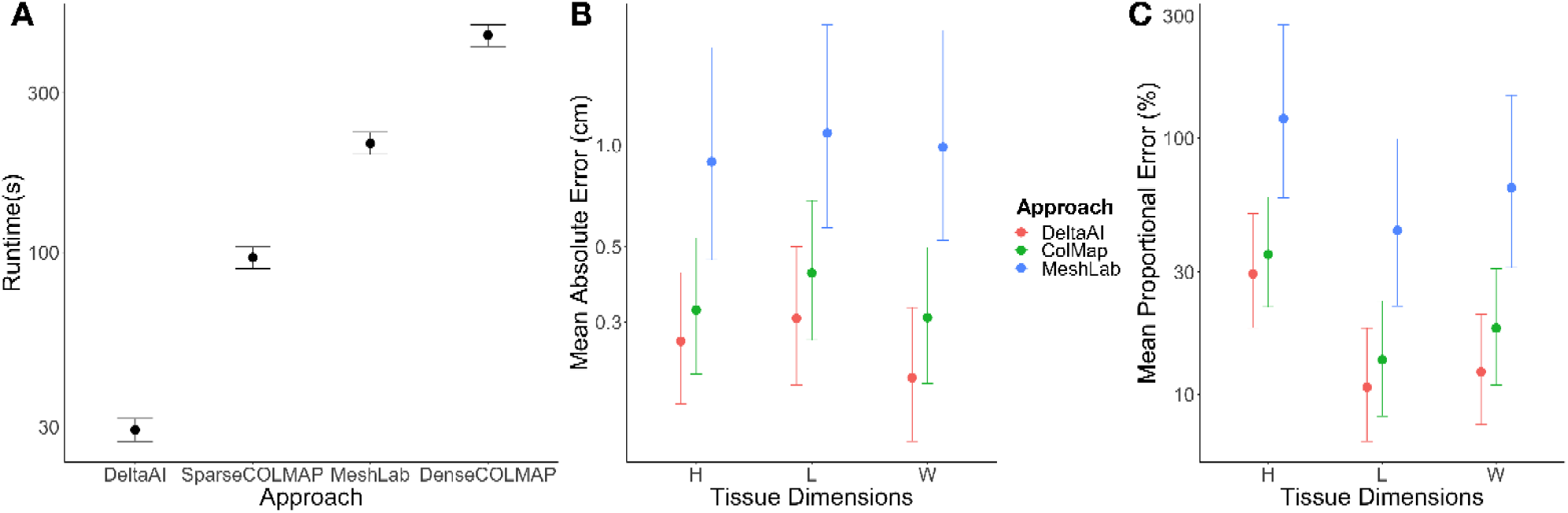
Model performance results– Average posterior estimates and posterior credible intervals comparing 3D modeling approaches for. **A)** Runtime, reported in seconds, **B)** Mean absolute error (MAE), reported in centimeters, and **C)** Mean proportional error, reported as a percentage of the original tissue dimensions

**Table 1:**
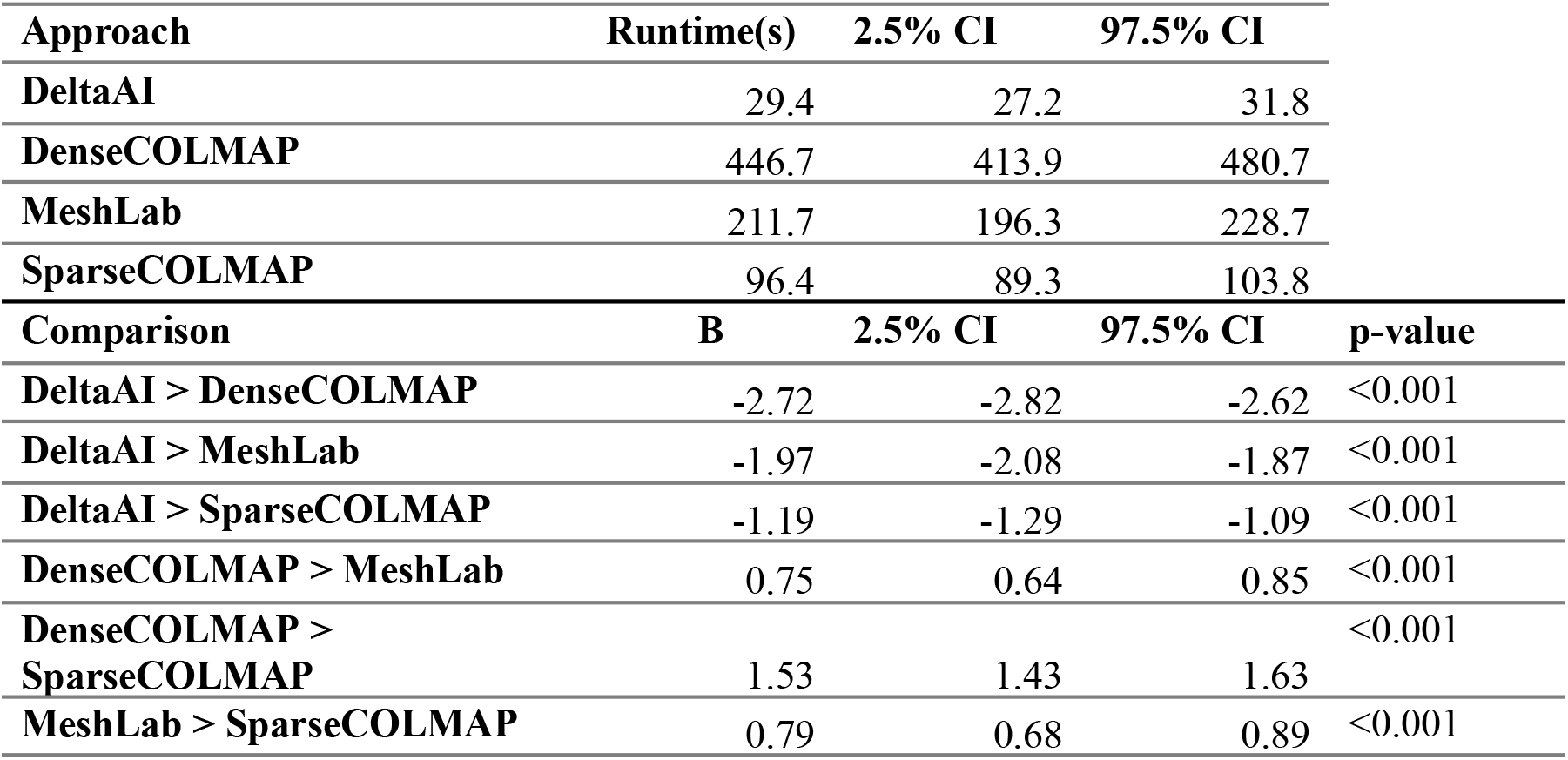
Average runtime for each approach and statistical comparison between runtimes. 95% credible interval estimates estimated by sampling the posterior distribution; each comparison tests to see whether runtime for one approach is higher than another; a negative coefficient (B) indicates that runtime of the first approach is less than the second approach.

| Approach | Runtime(s) | 2.5% CI | 97.5% CI |  |
| --- | --- | --- | --- | --- |
| DeltaAI | 29.4 | 27.2 | 31.8 |  |
| DenseCOLMAP | 446.7 | 413.9 | 480.7 |  |
| MeshLab | 211.7 | 196.3 | 228.7 |  |
| SparseCOLMAP | 96.4 | 89.3 | 103.8 |  |
| Comparison | B | 2.5% CI | 97.5% CI | p-value |
| DeltaAI > DenseCOLMAP | -2.72 | -2.82 | -2.62 | <0.001 |
| DeltaAI > MeshLab | -1.97 | -2.08 | -1.87 | <0.001 |
| DeltaAI > SparseCOLMAP | -1.19 | -1.29 | -1.09 | <0.001 |
| DenseCOLMAP > MeshLab | 0.75 | 0.64 | 0.85 | <0.001 |
| DenseCOLMAP > SparseCOLMAP | 1.53 | 1.43 | 1.63 | <0.001 |
| MeshLab > SparseCOLMAP | 0.79 | 0.68 | 0.89 | <0.001 |

### Measurement Results

The average of the length, width, and height MAE values for the tissue model generated by DeltaAI and other approaches are shown in **Figure 2B** and **Supplementary Table 2**. While DeltaAI provided more accurate measurements than other approaches (Z=4.43, p<0.001, **Table 2**), similar trends were found for proportional differences between predicted and measured tissue dimensions (**Figure 2C; Supplementary Table 3**). MeshLab was unable to converge on 58% of the tissue input data due to its inability to perform sufficient feature matching.

**Table 2:** Mean absolute error for each approach by tissue dimension and statistical comparison between runtimes. 95% credible interval estimates estimated by sampling the posterior distribution; each comparison tests to see whether mean absolute error for one approach is higher than another; a negative coefficient (B) indicates that mean absolute error of the first approach is less than the second approach; COLMAP results are reported for the best performing approach between sparse and dense reconstructions

Panel A
| Approach | Tissue Dimension | Mean Absolute Error (cm) | 2.5%CI | 97.5%CI |
| --- | --- | --- | --- | --- |
| COLMAP | H | 0.33 | 0.21 | 0.53 |
| DeltaAI | H | 0.26 | 0.17 | 0.42 |
| MeshLab | H | 0.89 | 0.46 | 1.94 |
| COLMAP | L | 0.42 | 0.27 | 0.69 |
| DeltaAI | L | 0.31 | 0.20 | 0.50 |
| MeshLab | L | 1.09 | 0.57 | 2.27 |
| COLMAP | W | 0.31 | 0.20 | 0.50 |
| DeltaAI | W | 0.21 | 0.13 | 0.33 |
| MeshLab | W | 0.99 | 0.52 | 2.18 |

Panel B
| Comparison | Tissue Dimension | B | 2.5%CI | 97.5%CI | p |
| --- | --- | --- | --- | --- | --- |
| COLMAP > DeltaAI | H | 0.21 | -0.37 | 0.81 | 0.462 |
| COLMAP > MeshLab | H | -1.01 | -1.84 | -0.25 | 0.009 |
| DeltaAI > MeshLab | H | -1.22 | -2.07 | -0.46 | 0.001 |
| COLMAP > DeltaAI | L | 0.31 | -0.29 | 0.88 | 0.308 |
| COLMAP > MeshLab | L | -0.95 | -1.80 | -0.19 | 0.016 |
| DeltaAI > MeshLab | L | -1.27 | -2.09 | -0.48 | 0.002 |
| **COLMAP > DeltaAI** | **W** | 0.41 | -0.17 | 1.01 | 0.169 |
| **COLMAP > MeshLab** | **W** | -1.16 | -2.05 | -0.38 | 0.002 |
| **DeltaAI > MeshLab** | **W** | -1.59 | -2.41 | -0.82 | <0.001 |
| **DeltaAI improves on COLMAP** | **Overall, across L/W/H** | Z=4.43 |  |  | <0.001 |

### Visualization of Modeling Results, Automated Tissue Grossing Measurements, and Placement of Tissue Orientation/Grossing Indicators

DeltaAI clearly produced the most visually consistent and high-quality representations of tissue compared to COLMAP and MeshLab (**Figure 3**). The sparse and dense COLMAP implementations required around one and a half minutes and ten minutes to compute a sparse and dense point cloud representation of the input tissue samples respectively (Levy et al., 2022). The point clouds generated using the COLMAP 3D tissue reconstruction revealed several holes and inconsistencies and were of lower resolution (similar to MeshLab). MeshLab produced models with extensive color inconsistencies, holes, and geometric inaccuracies. We were able to place orientation indicators (i.e., red and blue lines) and grossing indicators suggesting where to bisect tissue for further sectioning with ease for all three approaches (**Figure 4**).

**Figure 3:**
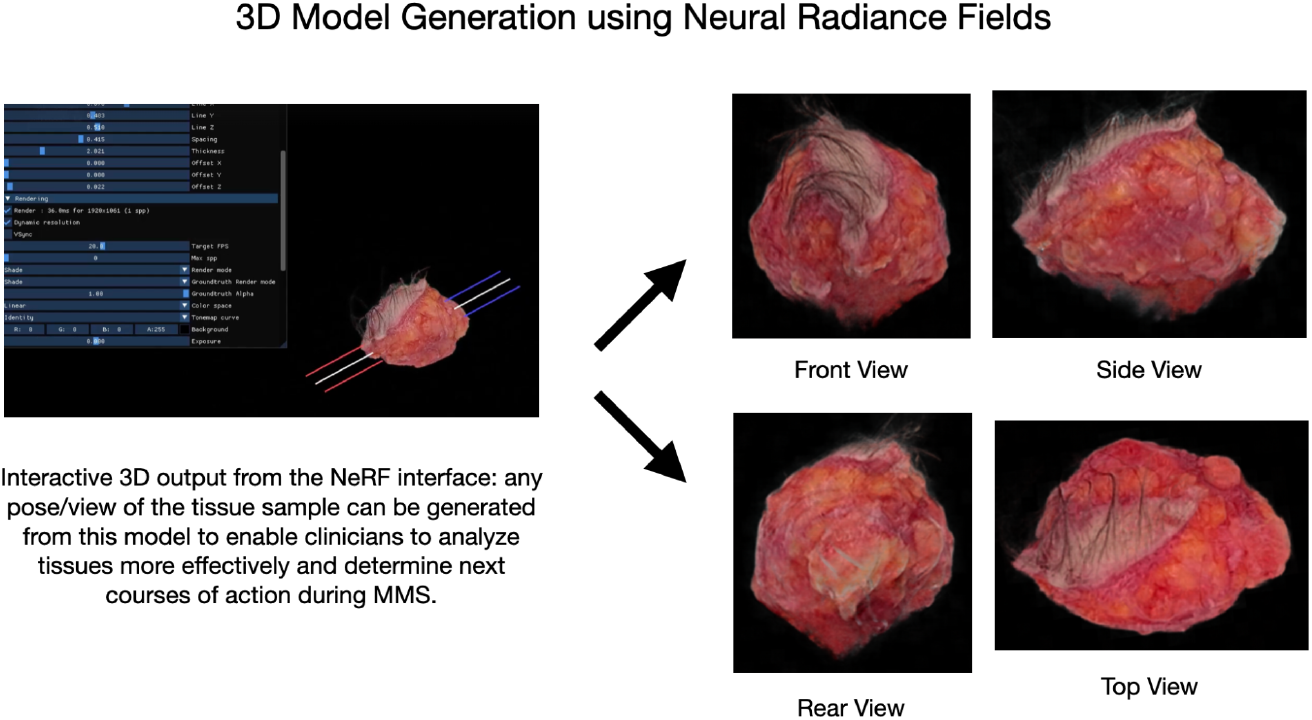
Three hundred sixty-degree “multi-directional” view of the tissue. The left side contains the output after training the NeRF model, which takes under 30 seconds to render the 3D model. The other side contains a portion of the various views that can be generated by using our 3D model’s interactive interface.

**Figure 4:**
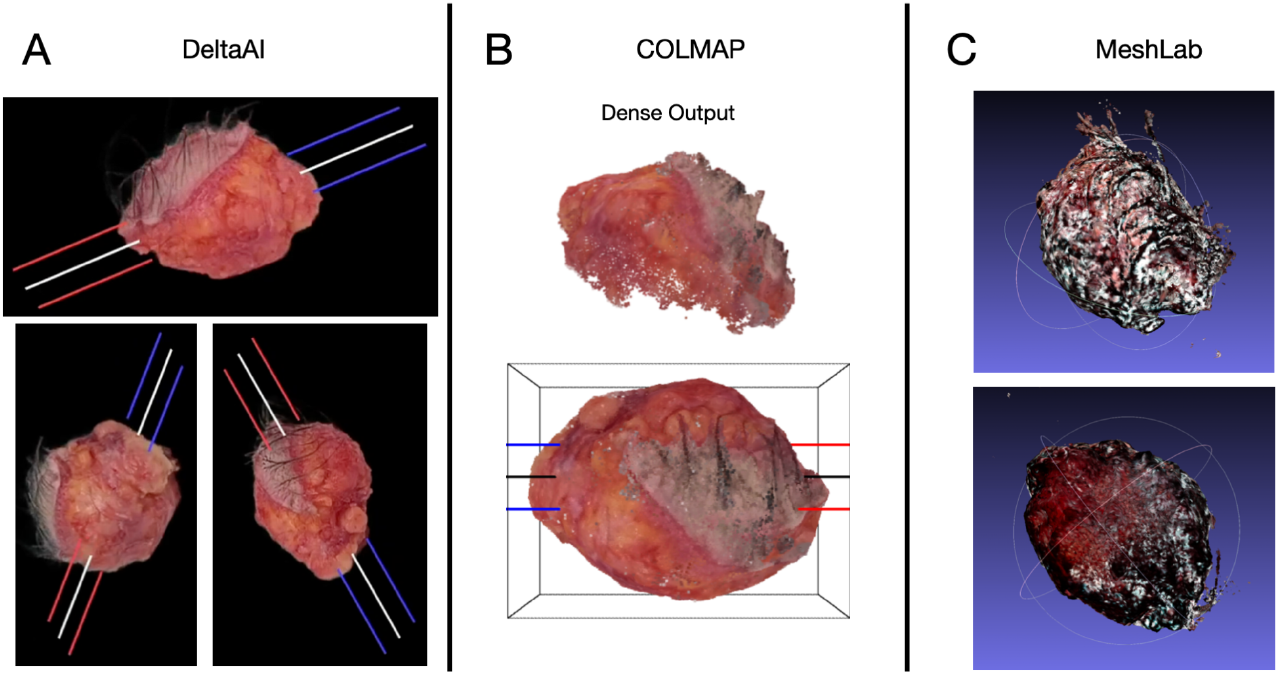
Visual comparisons of display outputs across approaches. **A)** The NeRF 3D model had superior results when compared to two existing photogrammetry models (MeshLab, COLMAP). More specifically, 3D tissue models produced from the NeRF model were faster and more accurate in terms of 3D model quality, measurement, and orientation. **B)** COLMAP is an existing workflow that performs point-cloud generation based on the triangulation of imaging features from multiple viewpoints, and the output 3D model of the tissue sample is shown, including the sparse (only one example shown) and dense point cloud (combining multiple sparse reconstructions) formation. **C)** The same input data was also imported to MeshLab’s software workflow, which had severe discoloration and low accuracy.

**Figure 5:**
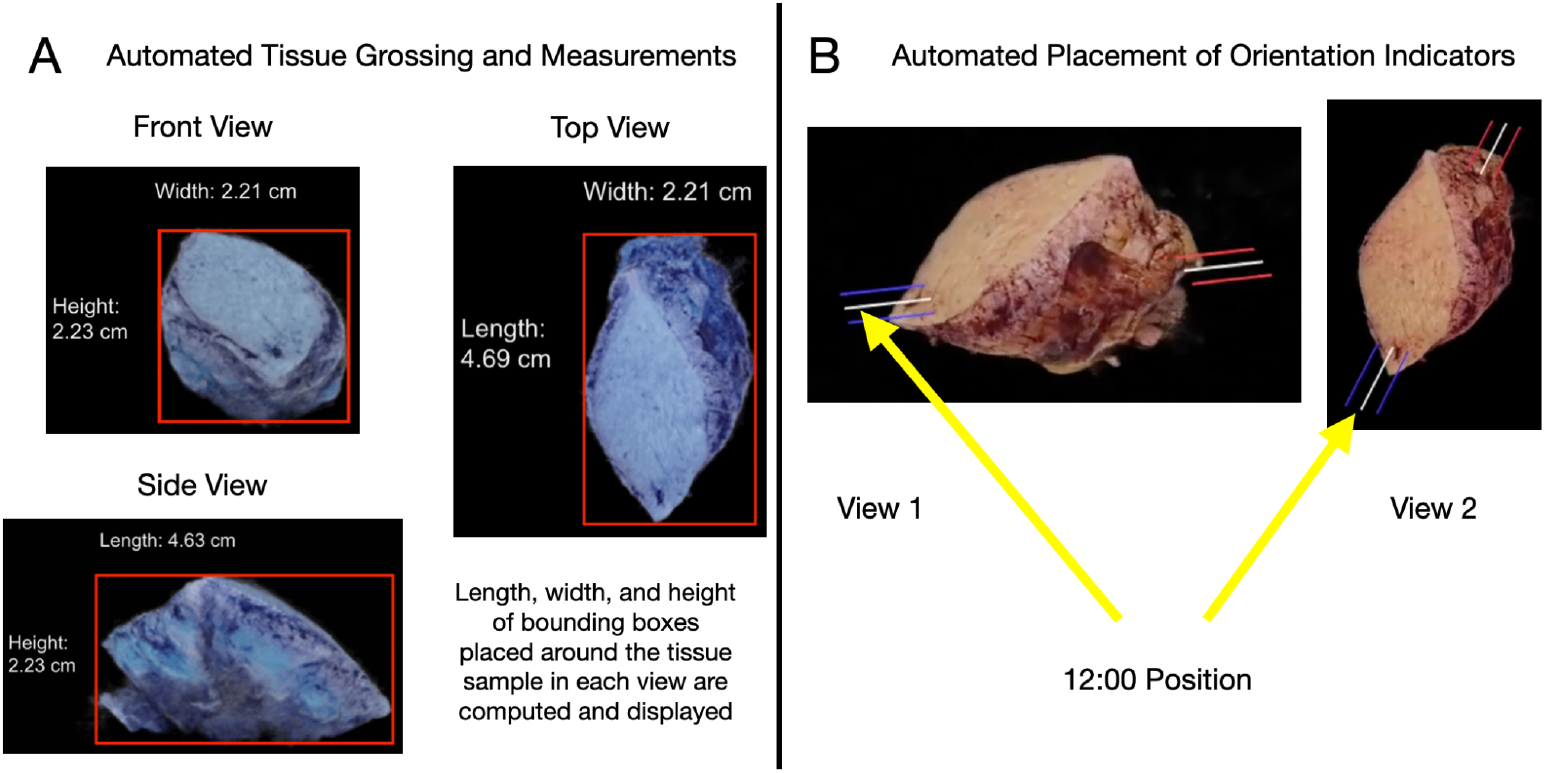
Measurement and Tissue Orientation/Grossing Indicators. **A)** This section contains the three separate views for the automated tissue grossing and measurement analysis. After placing the bounding boxes around the tissues using our object detection algorithm, the tissue dimensions were predicted by inputting camera parameters corresponding to front, top, and side views for the automated transfer of the inferred tissue dimensions to the EHR. **B)** This section contains two separate views for the orientation indicators. A total of three lines are placed through the tissue on both sides, indicating the 12:00 (blue) and 6:00 (red) positions and creating a visible plane (white) for bisecting the tissue.

## Discussion

Automating grossing measurements of a tissue sample during the pathology process can provide information about the size and shape of the tissue sample, which is important for certain types of tumors or other abnormalities as well as providing further means to optimize tissue sectioning (Beaulieu et al., 2018). For example, depth of invasion is an important metric for many cancers, especially cutaneous melanoma. When the depth of invasion data is paired with information on tissue weight and the histological assessment results (e.g., size of the tumor), it can be used to estimate tissue sample and tumor volume or weight, and the amount of tissue that is available for further analysis (Beaulieu et al., 2018). Accurate assessment of available tissue is crucial for ensuring that there is sufficient tissue for all the necessary tests and procedures, such as staining and microscopy, improving laboratory automation which can expedite manual and or semi-autonomous histological slide interpretation. Finally, grossing measurements can help to standardize the process of tissue preparation and analysis, which is important for ensuring the accuracy and reproducibility of the results (Bialy et al., 2004). By taking consistent and accurate measurements, pathologists can ensure that they are working with tissue samples that are representative of the original tissue, which can help to improve the reliability of their findings (Bialy et al., 2004). Furthermore, the workflow established through Neural Radiance Fields enables this tissue grossing analysis to occur with precision and speed, freeing up a technician to perform other tasks.

NeRF’s dual-modality as a comprehensive renderer and a volumetric modeler coupled with its ease of setup makes it an impelling option for high-quality photogrammetry. Compared to COLMAP’s dense reconstruction to create a point cloud representation, the new model converged significantly faster without the holes or color inconsistencies that are typically representative of point cloud-based approaches. The success of NeRF is largely due to the architecture of Neural Radiance Fields, which enables the model to dynamically optimize the 3D reconstruction as it dynamically processes images filling a gap in existing artificial intelligence workflows and software that lack within the domain of intraoperative margin assessment (Jiang et al., 2022).

Applying NeRFs, DeltaAI convincingly demonstrated the ability to reconstruct entirely novel (i.e., unobserved) views, with only partial information about the view from each frame, while retaining volumetric information in the process. Even the various views of the tissue with little information present in each frame, such as the bottom view, still retain proper proportions despite being discolored. NeRF offers millisecond render rates, allowing for real-time rendering within a mobile application setup. However, for a few of the tissue samples, the NeRF model was not able to converge on a high-resolution, 3D output after rendering. The size, diversity, and quality of the input data influence the accuracy and efficiency of a NeRF model. Pre-training methods which form a gestalt impression of myriads of tissue shape characteristics prior to finetuning on a new sample may further improve results (Bisson-Larrivée & LeMoine, 2022). The scope and quality of input data in the training of NeRF models is critical in determining algorithm run time, i.e. time to complete a 3D rendering (Bisson-Larrivée & LeMoine, 2022). Increased size and quality of data, leads to increased processing power and time requirements. Despite this, it is essential to maintain the diversity and quality of inputs to allow for more broad generalization across all tumor types, sizes, and locations, regardless of unique tissue characteristics (Mastacouris & Mafee, 2021). The balance between dataset size and input image quality/diversity may determine workflow effectiveness (Sarker, 2021).

Pre-training techniques are used in several different machine-learning architectures and algorithms. They allow for a more accurate output or result given even a smaller dataset since the system has already been trained on other data (Han et al., 2021). Pre-training the NeRF model on a large dataset can also help improve its performance on a specific task providing it with a strong foundation of knowledge about the task at hand. This can help the sequence converge faster and achieve better performance in a more accurate and timely manner. In the future, this is a technique that should be integrated into all Neural Radiance Fields workflows. Oftentimes, for heavy-duty tasks, the time can take anywhere from 10-25 hours to complete, sometimes without convergence of the 3D model (Zhang et al., 2020). This massive amount of time is not feasible for this NeRF model, especially during MMS, where rapid grossing assessments and visualizations are required, meaning that integrating pre-training techniques is essential to this entire process, especially when transitioning to more complex or larger tumor types (e.g., head and neck squamous cell tumors) (Levy et al., 2022).

Our DeltaAI NeRF algorithm can be useful during Mohs surgery in a multitude of ways. First, our workflow can generate real-time 3D visualizations and depictions of the tissue during surgery. Combining 3D modeling results with histological findings can help surgeons better understand the spatial relationships between various structures found inside of the tissue or that are generally in that location, such as tumor relationship to margin, blood vessels, nerves, and other critical components, for further real-time treatment planning. Another factor is the improved accuracy conferred from the usage of Neural Radiance Field models. High-fidelity tissue representations allow histologists and surgeons to optimize tissue sectioning precisely identify and remove malignant tissue while limiting the damage to healthy tissue, improving recurrence, and functional and aesthetic outcomes. The third impact of our algorithms on MMS is the automation factor. NeRF approaches are highly automated (i.e., point and click, from video to model), reducing the time and effort spent on creating 3D models of the tissue for guiding specimen grossing and reporting of the tissue dimensions. Overall, this can help to improve the efficiency and rapidness of Mohs surgery and allow surgeons, nurses, and technicians to focus on other critical aspects of the procedure. The rendered 3D model can also be leveraged for post-operative analysis, providing a vivid and detailed, interactive representation of the tissue that was removed during the surgery *prior* to sectioning. This feature can provide tumor boards with a more in-depth understanding of the 3D involvement of the tumor and plan for future treatment while iteratively improving surgical techniques personalized to the patient’s characteristics.

Compared to conventional cancer resection methods, Mohs micrographic surgery (MMS) offers a real-time assessment of tumor margins, reducing the risk of tumor recurrence and preserving healthy tissue (Morman, 1989). However, the current MMS workflow is a multi-step process that involves multiple individuals, increasing the chances of error and adding stress and cognitive load during surgery (Ecclestone et al., 2020). In this study, we developed DeltaAI, a machine learning platform based on Neural Radiance Fields, which generated a 3D model of tissue samples in under 30 seconds, without any gaps, holes, or inconsistencies. This rapid and accurate analysis of the entire tissue sample could provide pathologists and histo-trained surgeons with more precise recommendations for further tumor removal steps. Additionally, the automated reporting of tissue dimensions and the placement of grossing recommendations indicators streamline documentation and reduce manual labor. By harnessing state-of-the-art deep learning 3D rendering systems, this tool has the potential to enhance operational efficiency and facilitate further workflow improvements. Future research will focus on developing an enhanced mobile application interface using augmented reality and a touch-based visualization system, building upon this original research. In conclusion, we have successfully developed a novel 3D tissue mapping and reconstruction technique for MMS tumor resections utilizing NeRF. Our model demonstrated superior performance in terms of speed, accuracy, and flexibility compared to existing models. This advancement could enable more automated and quantitative tissue processing and analysis, leading to improved outcomes in Mohs micrographic surgery.

## Supplementary Material

### Supplementary Methods

#### Data Collection and Study Population

In our study, we utilized photogrammetry techniques to create 3D models of tissue samples for histological assessments. This cost-effective approach involved using a phone camera placed at a fixed distance from a turntable. By recording low-resolution videos of tissue samples rotating on the turntable, we developed a tissue grossing algorithm and generated 3D models.

During Mohs surgery, we positioned 17 large tissue samples on circular turntables and recorded 5-second video clips of each rotation, resulting in 946 frames per video. To simulate faster-spinning tables for future setups, we sampled longer videos. These videos were then processed using DeltaAI, a machine-learning model, to generate sophisticated 3D tissue reconstructions. The size measurements of the tissue samples were obtained using an automated calibration tool.

#### Neural Radiance Fields

Neural Radiance Fields (NeRF) is a machine learning architecture used for generating 3D models based on input images from different camera positions (Zhang et al., 2020). This architecture employs radiance fields to describe the light emitted or reflected by objects in space (Goel et al., 2022). Radiance fields consider lighting conditions, material properties, and surface geometry, making them useful for generating realistic images and reconstructing scenes (Goel et al., 2022).

To create and train a NeRF model, images of a three-dimensional structure and corresponding camera poses are required (Zhang et al., 2020). The model learns the features and characteristics of the object based on this data, constructing a densified 3D structure aligned with the input images. This results in an interactive 3D rendering showcasing different perspectives of the object (Zhang et al., 2020). Neural Radiance Fields have diverse applications, enabling animations of 3D objects and real-time visualization (Gao et al., 2022).

In 3D visualization and rendering, a pose represents the position and orientation of an object or camera in 3D space. It can be represented by translation and rotation parameters, facilitating the transformation between coordinate frames for generating a 3D visualization/render. Intrinsic parameters describe the internal characteristics of the camera, such as focal length and distortion, while extrinsic parameters specify the camera’s orientation and location in 3D space. Together, these parameters allow for modeling the transformation between the 3D world and the 2D image captured by the camera. Understanding camera intrinsics and extrinsics is crucial for automated 3D tissue reconstruction (Bisson-Larrivée & LeMoine, 2022).

The applications of NeRF include portable 3D-mapping systems that use RGB cameras and inertial measurement units (Farina et al., 2022). Advances in training and rendering techniques have significantly reduced the convergence time of NeRF models, enabling faster results in minutes or even seconds (Müller et al., 2021).

#### DeltaAI: Model Architecture

Our novel implementation of machine learning utilizes a NeRF model trained with hash encodings that allow for rapid convergence. This 3D model can utilize machine learning to automatically render and fill in gaps/holes in the tissue sample (something other techniques are unable to accomplish) (Müller et al., 2021). Its capacity to create intricate 3D models from a small collection of 2D photos is why this model was selected. NeRF learns a continuous function describing the 3D scene from a collection of 2D input photos with the associated camera postures. The primary novelty of the model is the use of a revolutionary sampling technique to drastically lower NeRF’s computational complexity, enabling real-time rendering of 3D scenes on low-cost hardware.

Encoders and decoders make up the architecture of the Instant-NGP NeRF model. A latent space representation is created by the encoder using the camera posture and several 2D picture attributes as input. The decoder converts a 3D point’s RGB color and opacity value from the point’s input, which consists of a point in space, into the camera posture. The volumetric rendering loss, which evaluates the discrepancy between the predicted color and opacity and the ground truth color and opacity of each point in the 3D space, is used to train the decoder.

The model developed in this study, DeltaAI, has a significant advantage over conventional 3D reconstruction techniques, which need a large number of photos and intricate calibration procedures to produce highly detailed 3D models from a sparse set of 2D images. Even in the face of noise and occlusions, our model’s machine learning component allows it to understand the underlying structure of the image and produce precise 3D representations. The 3D reconstruction of tissue samples using DeltaAI was effective and precise, which is essential for many biological applications.

#### Preprocessing

During the tissue sample data collection process, a 360-degree capture was taken using a roundtable setup. However, the images obtained had to be processed to eliminate the background objects and retain only the tissue sample. For this purpose, a segmentation model was employed to localize and crop the tissue sample. Each pixel in the image is examined by the segmentation model, which groups them into objects based on how similar their colors, textures, and other characteristics are. After the desired criteria have been defined, the model removes the objects that do not match them. In our scenario, the model was trained to distinguish the tissue sample from any background objects based on its distinctive characteristics, such as form, size, and color. We made sure that the input data for the 3D reconstruction method only included the tissue sample by utilizing the segmentation model to exclude any other items that would have hampered the model’s convergence. This produced an internal 360-degree capture of the tissue sample, where photos were recorded from various perspectives and angles, giving the impression that the camera was spinning while in fact, the roundtable was doing the spinning. Camera frames were taken strategically at intervals to ensure the highest performance and efficiency of the data inputted into the NeRF model.

During capturing the tissue samples, we observed that motion blur was a notable issue at the start and end of the videos. To address this issue, we employed a sharpness model to identify and remove blurry frames from the dataset. The sharpness model is a computational algorithm that computes the sharpness or focus of an image by analyzing various image features such as edges, texture, and contrast. The sharpness model then applies a threshold value to the computed sharpness measure, below which an image is considered blurry and subsequently removed from the dataset. By using the sharpness model to weed out blurry frames, we were able to ensure that the resulting dataset consisted only of clear and focused images, which were suitable for subsequent processing and analysis.

We found that an image set of 20-40 images allowed our model, DeltaAI, to converge the fastest, which can be acquired using a turntable that can run in X-second revolutions. The estimation of poses is a crucial step in training any neural radiance field model. In our work, we employed COLMAP, a library, and platform created by Johannes L. Schönberger and Jan-Michael Frahm, to estimate the poses. COLMAP is capable of performing full 3D reconstruction and photogrammetry using a set of 2D images. To optimize the performance of the pose estimation process, we utilized a sparse reconstruction approach. Specifically, we used a third of the available image set to estimate poses and construct the sparse model. The remaining poses were subsequently filled in while holding the roundtable and camera motion constant. The sparse reconstruction approach enables the determination of poses in under 10 seconds, since only a few 3D points need to be triangulated, making it a highly efficient and effective method. Notably, the use of a sparse model suffices for this task, given that only the estimation of poses was required, and the resulting model from COLMAP does not need to be highly detailed, due to NeRF’s ability to render these high-resolution models.

Camera intrinsics from an iPhone 12 (Apple, Cupertino California) were used to calibrate the camera model, while translation and rotation (extrinsic) were estimated by COLMAP. This data was all inputted into the NeRF model, which automatically filtered out certain input images with detrimental characteristics, including low resolution, blur, and lack of contrast, allowing for the model to avoid inconsistencies in the 3D reconstruction. After around 15 seconds, the loss began to plateau out, and the model converges successfully on a 3D visualization of the tissue. This 3D model is displayed in a graphical user interface, powered by the instant-ngp software, to render the views of the tissue after it has been triangulated, and perform further grossing measurements and analysis, as well as assess the NeRF model performance.

### Comparison Methods

#### MeshLab and COLMAP

MeshLab is an unrestricted and fully-accessible programs and can perform photogrammetry, which is the process of creating a 3D visualization from a set of 2D images (Cignoni et al., 2008). MeshLab was used as a base comparison to assess the performance of the NeRF workflow that we developed and determine its advantages relative to popular photogrammetry techniques. COLMAP is an open-source software that is commonly used for 3D reconstruction, camera localization, and image-based rendering. Although COLMAP’s ability to perform full 3D reconstruction is limited, its automated pose estimation feature is highly effective. We utilized this aspect of COLMAP’s workflow in our DeltaAI platform, which is based on Neural Radiance Fields (NeRF) to accurately estimate the camera orientation and position for each image in our dataset. While we did not use COLMAP’s 3D reconstruction process in DeltaAI due to its inefficiency, we used it to compare our model’s performance with an existing model. This allowed us to validate the accuracy of our models and make necessary adjustments to further optimize our platform.

#### Automated Tissue Grossing Measurements

Automated grossing measurements including the length, width, and height of the tissue sample are automatically calculated using a sophisticated object detection algorithm that places bounding boxes over the tumors to identify major axes, and subsequently, tissue dimensions. Effectively, six views of the tissue are generated, including anterior, posterior, superior, inferior, and lateral views. From these perspectives, the dimensions of the tumor were calculated and then averaged among the corresponding axis of each view to allow for the highest accuracy. These measurements were determined by the specific RGB values and a contour detection algorithm to place a rectangular bounding box around the tissue. The pixel coordinates were later converted to centimeters using a calibrated conversion scale, providing automated and accurate representations of the tissue size. This conversion scale was formed by placing a fiducial ellipsis of known size on the turntable in the input dataset, which provided the unit conversion from pixels to real-world centimeters and efficient comparison to assess the measurement model performance.

#### Orientation Tissue Indicators

Orientation indicators were mapped to the 12:00 position (superior) of the tissue sample and run through the body of the 3D tissue model, aligning properly from both ends. As this position is difficult to estimate from the 2D image set using only the angle at which the tissue is acquainted, the COLMAP sparse reconstruction was used instead. This 3000-5000 point cloud construction was sufficient to determine the 12:00 position of the tissue and draw a line from the centroid of the tissue sample to the 12:00 position. The same line is flipped to the other side of the tissue to define the opposite pole (inferior) of the tissue as well. The position of the line is then mapped onto the original image set and the NeRF model is retrained with the lines, causing the indicator lines to automatically show up in the reconstructed NeRF model in DeltaAI.

This methodology and workflow can be used for any number of indicators or annotations required, and for our tissue samples, we have generated three different lines which are oriented in the direction of the 12:00 position. These three lines create a plane through the tissue sample inking points, which are generally near the center, allowing for the specific cutting points to be marked by the surgeons with ease. This process for automatically placing orientation indicators on the tissues is very important during Mohs Micrographic Surgery since it allows the surgeons and histologists to accurately map the tissue positions and sample additional tissue judiciously (Dinehart & Pollack, 1989). Additionally, this algorithm has a short runtime and is integrated with the Neural Radiance Fields model, allowing for the output 3D model to have orientation indicators placed on the proper positions without any extra manual work after the model has converged.

DeltaAI has the potential to work over several tissue sections in addition to the orientation tissue indications to offer a thorough 3D image of the tissue sample. A comprehensive 3D representation of the entire specimen may be created using our graphical user interface by smoothly integrating various tissue sections. By enabling the examination of the complete tissue margin, this feature lowers the possibility of missing any remaining cancer cells and the chance of recurrence. The GUI’s user-friendly platform, which will eventually be converted into a mobile application as well, enables real-time assessment of the tissue margins, effective surgical planning, and faster diagnosis. It can be seamlessly handled by doctors and other medical professionals. An essential tool in Mohs Micrographic Surgery and other surgical procedures, DeltaAI’s capacity to work across many tissue sections and produce a whole 3D model of the tissue sample makes it so.

## Supplementary Figures/Tables

**Supplementary Figure 1:**
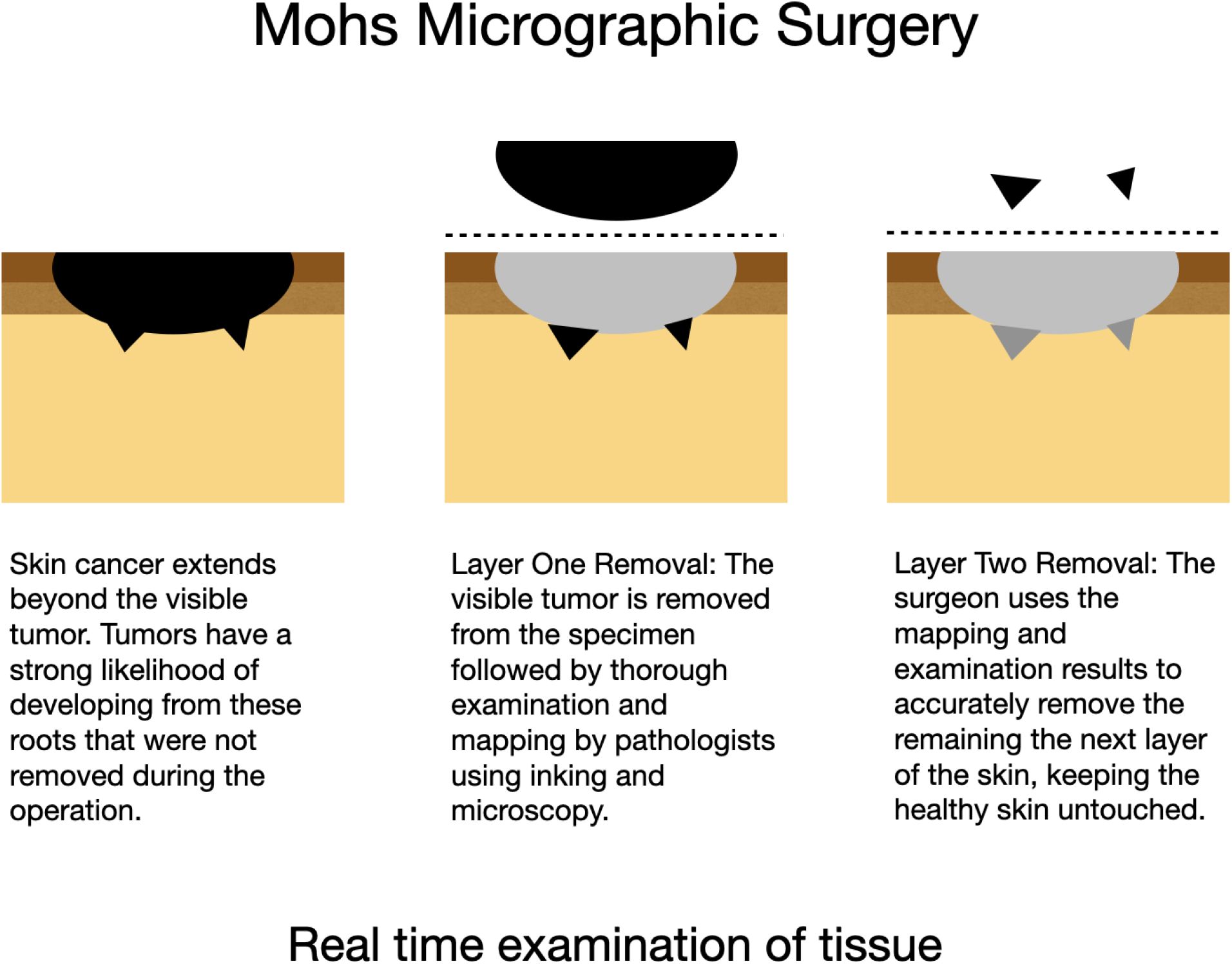
Mohs Micrographic Surgery Workflow. 1) a visible layer of the tumor is removed by the surgeon, 2) grossed (cryo-frozen, bisected where appropriate, applications of ink, sectioned, stained), 3) thoroughly examined by the histologists (or surgeon) through microscopy, 4) reporting results with respect to original anatomic position and orientation for 5) additional removal of remaining cancerous skin layers until 100% tumor removal.

**Supplementary Figure 2:**
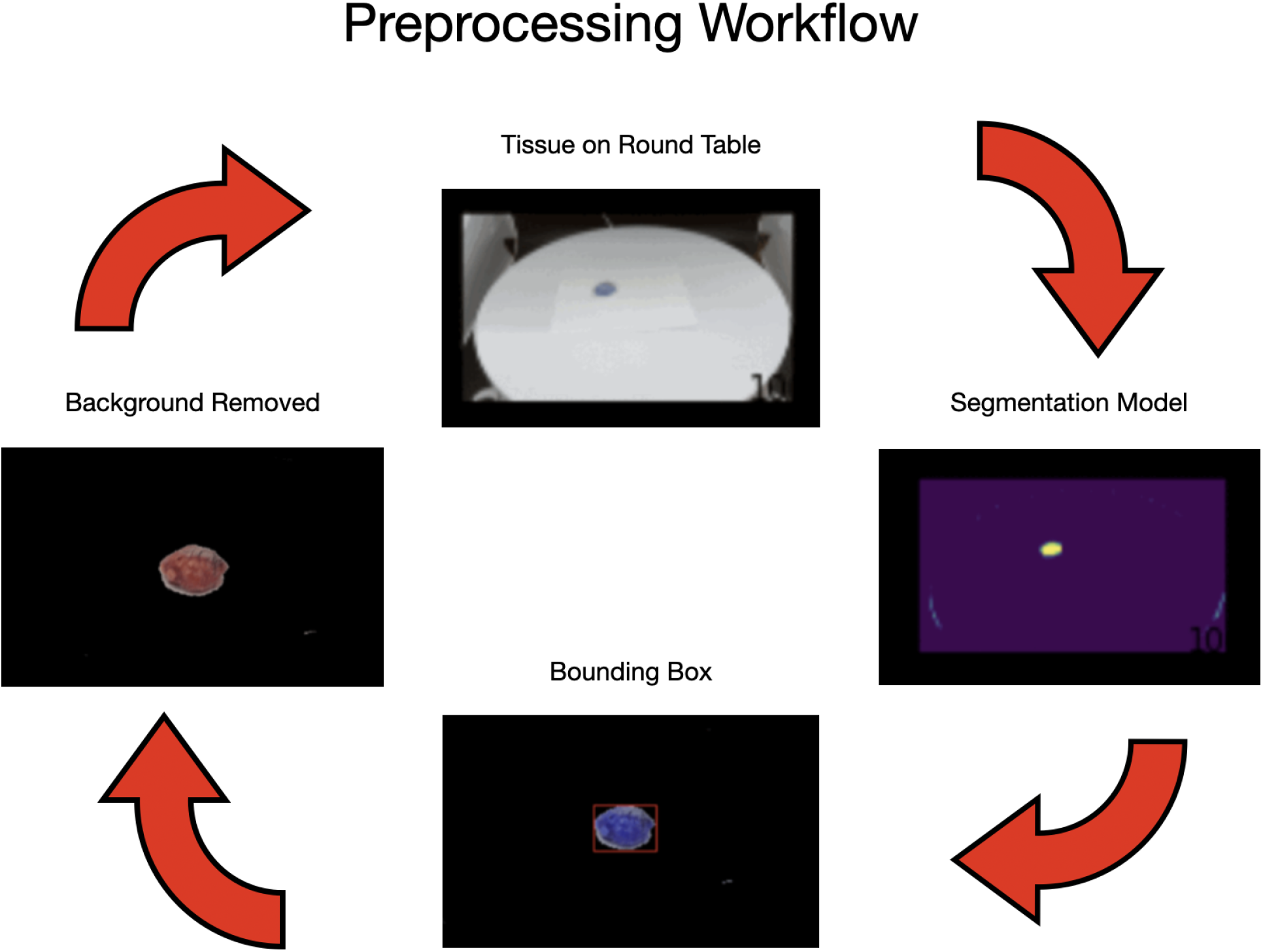
Preprocessing workflow overview. 1) video of issue is taken as it rotates on a roundtable, 2) Tissue samples were localized using a segmentation network, with the tissue emphasized in yellow, 3) bounding boxes were placed around the localized object and 4) the background was removed to prepare the data for the NeRF model.

**Supplementary Table 1:**
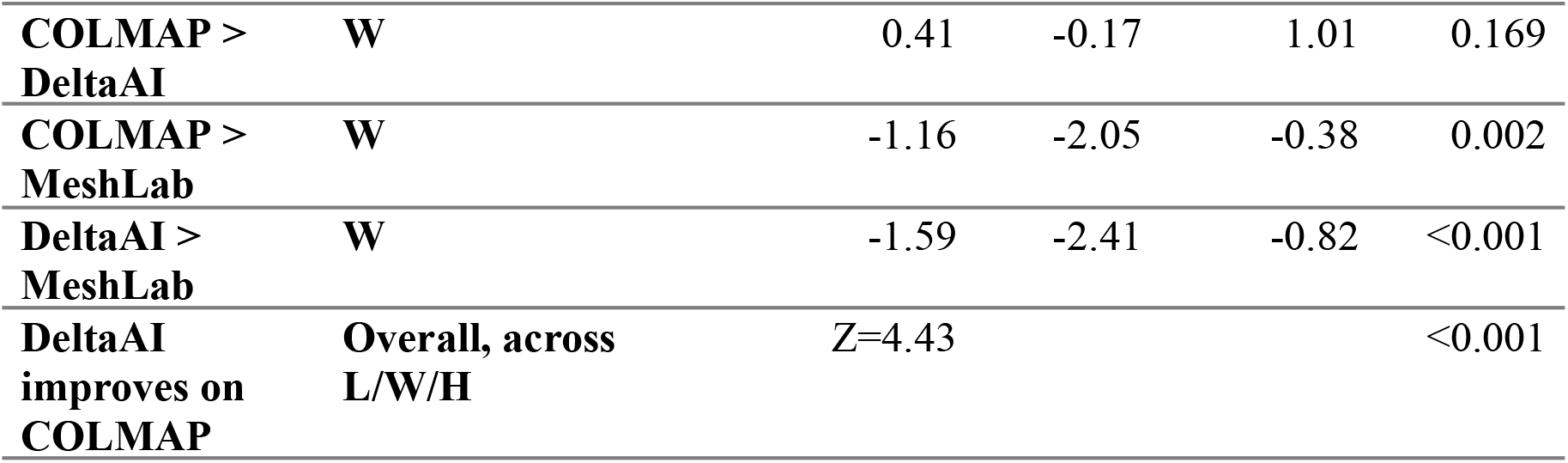
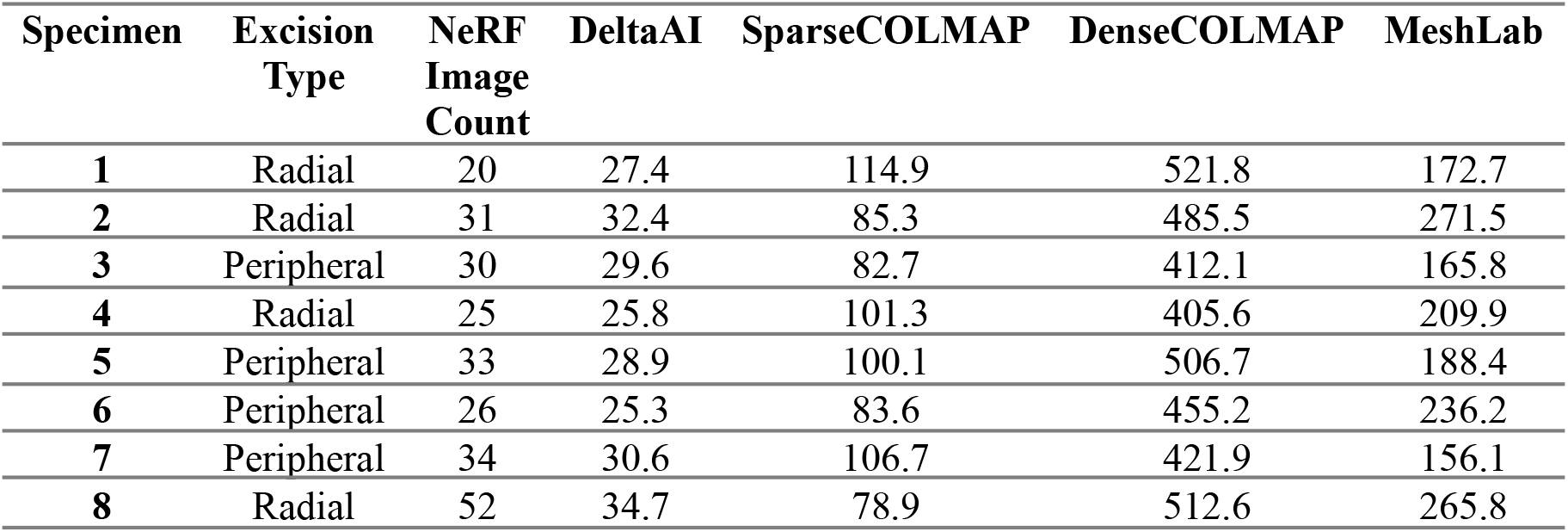
Contains runtime results for performing 3D modeling (pose/camera estimation and model training) for all three of the comparison methods (DeltaAI, COLMAP, MeshLab) for each specimen

**Supplementary Table 2:** Tissue dimensions as measured by the Mohs surgical staff, then using: COLMAP, DeltaAI and MeshLab, reported in centimeters; N/A indicates instances where MeshLab was unable to converge

| Specimen | Measured (cm) |  |  | COLMAP - Predicted (cm) |  |  |
| --- | --- | --- | --- | --- | --- | --- |
|  | L | W | H | L | W | H |
| **1** | 2.4 | 1.3 | 0.6 | 2.38 | 1.23 | 0.79 |
| **2** | 4.5 | 4 | 3.3 | 4.86 | 3.61 | 2.61 |
| **3** | 2.8 | 2 | 2.2 | 2.43 | 2.17 | 1.73 |
| **4** | 4.5 | 2 | 2.5 | 4.64 | 2.52 | 2 |
| **5** | 4.5 | 3.5 | 3 | 4.87 | 3.49 | 2.98 |
| **6** | 2 | 2 | 0.6 | 2.02 | 1.98 | 1.05 |
| **7** | 3.5 | 2.5 | 2 | 3.97 | 3.3 | 1.9 |
| **8** | 2 | 1 | 0.7 | 1.68 | 0.88 | 0.45 |
| **9** | 3 | 1.5 | 0.5 | 3.84 | 1.98 | 0.78 |
| **10** | 3 | 1.5 | 0.5 | 2.47 | 1.27 | 0.85 |
| **11** | 1.5 | 1.3 | 0.7 | 1.54 | 1.28 | 0.8 |
| **12** | 2.9 | 2.4 | 1.1 | 2.86 | 2.2 | 1.09 |
| **13** | 2 | 1.5 | 0.6 | 2.23 | 2.06 | 1.14 |
| **14** | 5.1 | 1.7 | 0.6 | 3.46 | 1.26 | 0.79 |
| **15** | 2.8 | 1.3 | 0.7 | 3.96 | 2.03 | 1.34 |
| **16** | 1.7 | 1 | 0.5 | 1.91 | 1.13 | 0.79 |
| **17** | 2.1 | 1.3 | 1 | 3.59 | 2.05 | 1.53 |
| Specimen | DeltaAI - Predicted (cm) |  |  | MeshLab - Predicted (cm) |  |  |
|  | L | W | H | L | W | H |
| **1** | 2.35 | 1.32 | 0.63 | 2.58 | 2.15 | 1.59 |
| **2** | 4.43 | 3.89 | 3.49 | N/A | N/A | N/A |
| **3** | 2.61 | 2.24 | 2.16 | 1.43 | 1.2 | 1.29 |
| **4** | 4.69 | 2.21 | 2.23 | 3.1 | 2.66 | 2.19 |
| **5** | 3.97 | 3.33 | 3.37 | N/A | N/A | N/A |
| **6** | 2.33 | 2.29 | 0.96 | N/A | N/A | N/A |
| **7** | 3.39 | 2.57 | 2.13 | N/A | N/A | N/A |
| **8** | 2.09 | 0.81 | 0.95 | 3.34 | 0.91 | 0.9 |
| **9** | 3.43 | 1.67 | 0.89 | N/A | N/A | N/A |
| **10** | 3.9 | 1.71 | 0.49 | N/A | N/A | N/A |
| **11** | 1.33 | 1.52 | 1.1 | 1.49 | 0.65 | 0.38 |
| **12** | 3.14 | 2.2 | 0.93 | 1.17 | 0.88 | 0.7 |
| **13** | 1.89 | 1.36 | 0.95 | N/A | N/A | N/A |
| **14** | 4.4 | 1.51 | 0.86 | 4.68 | 2.76 | 2.94 |
| **15** | 3.58 | 1.67 | 1.2 | N/A | N/A | N/A |
| **16** | 1.49 | 0.82 | 0.81 | N/A | N/A | N/A |
| **17** | 2.65 | 1.83 | 1.34 | N/A | N/A | N/A |

**Supplementary Table 3:**
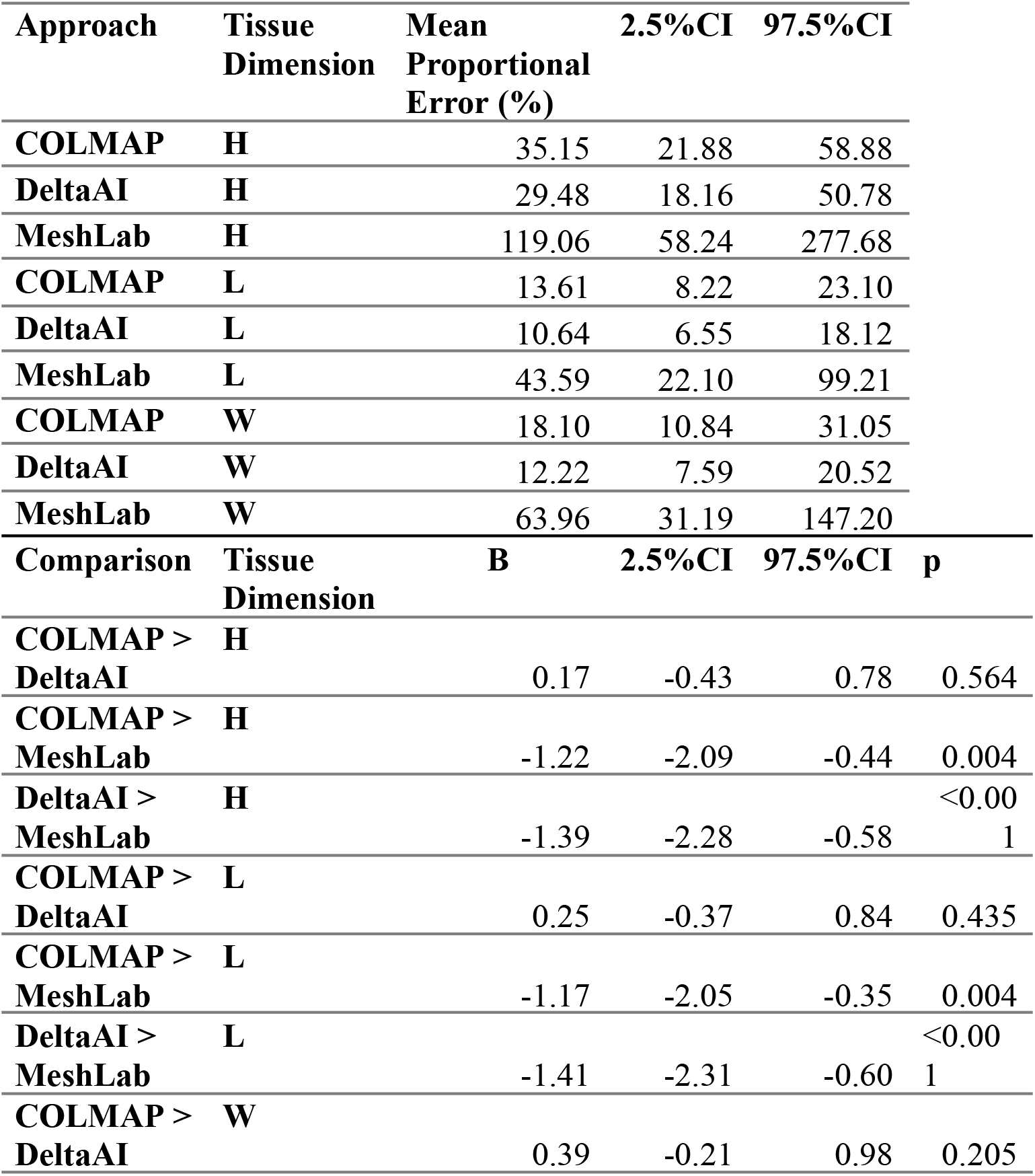
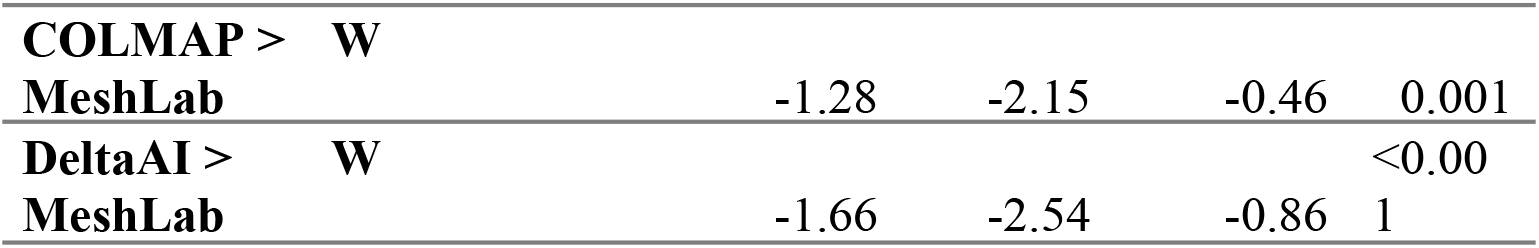
Mean proportional error for each approach by tissue dimension and statistical comparison between runtimes. 95% credible interval estimates estimated by sampling the posterior distribution; each comparison tests to see whether mean proportional error (%) for one approach is higher than another; a negative coefficient (B) indicates that mean absolute error of the first approach is less than the second approach; COLMAP results are reported for the best performing approach between sparse and dense reconstructions

**Supplementary Figure 3:**
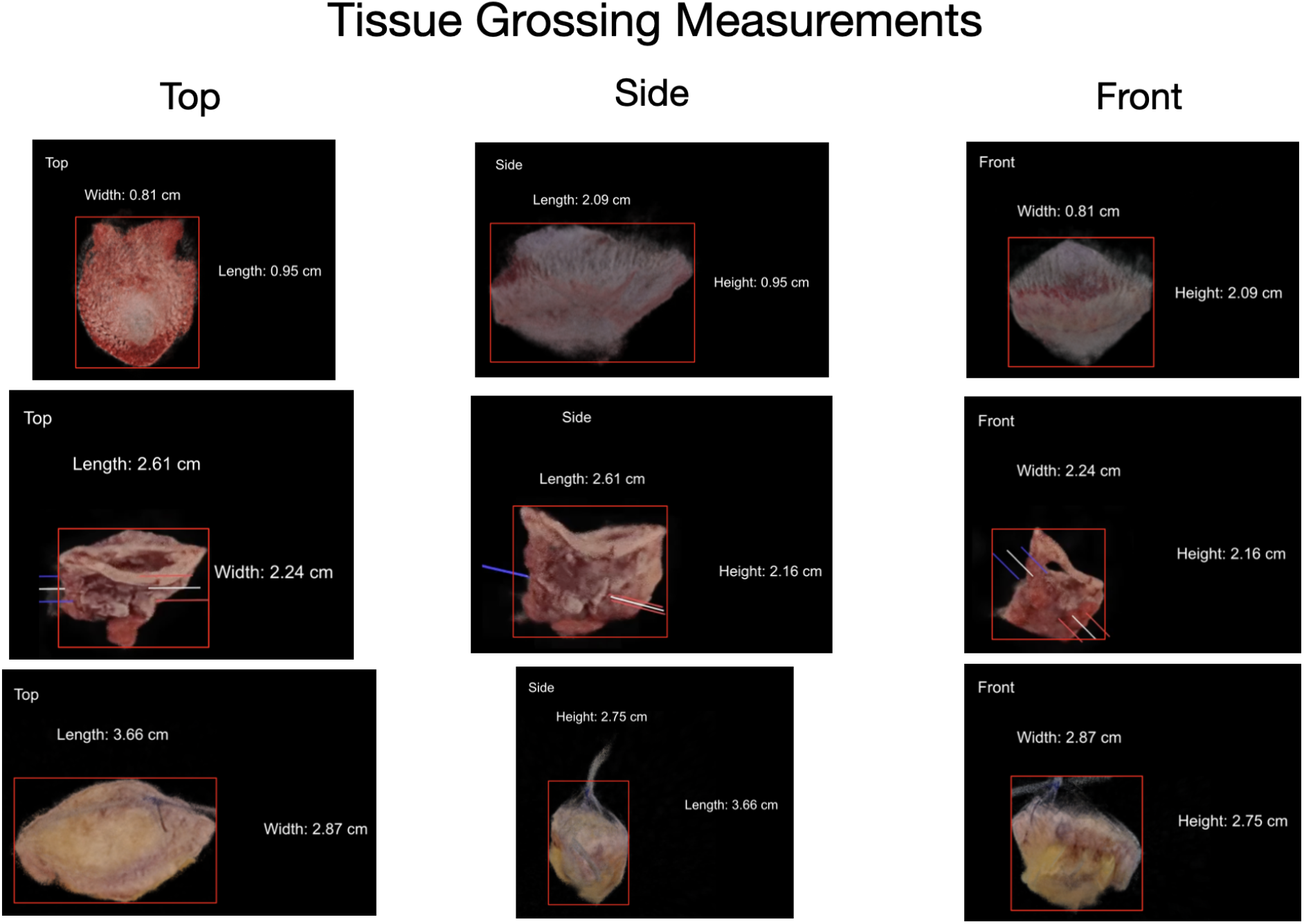
NeRF Results demonstrated with the tissue grossing measurements for three MMS-style specimen. These samples were grossed using the 3D rendering output from the DeltaAI workflow, along with the volumetric representations generated which made it seamless to estimate tissue size.

